# Dispelling the Plateau Assumption: Case-Based Evidence for Language Recovery in Chronic-Post Stroke Aphasia

**DOI:** 10.64898/2026.10.03.755714

**Authors:** Elizabeth H.T. Chang, Alycia B. Laks, Kyle Shattuck, Andrew T. DeMarco, Peter E. Turkeltaub

## Abstract

**Objective:** Recovery from post-stroke aphasia is traditionally thought to plateau within a year after stroke. As such, no studies have characterized long-term recovery using comprehensive behavioral assessment and longitudinal functional MRI. We sought to characterize behavioral and neuroplastic changes over nearly six years in a patient with chronic post-stroke aphasia.

**Methods:** A patient with chronic left hemisphere stroke underwent behavioral assessment and task-based fMRI at 9, 22, 33 and 71 months after stroke. Testing assessed auditory comprehension, semantic and phonological processing, naming, reading, spelling, motor speech, and executive function. Task-related activation during an adaptive semantic matching paradigm was quantified within language and semantic regions of interest.

**Results:** Behavior improved across multiple language domains. WAB-R Aphasia Quotient increased from 49.2 to 69.2, along with the large gains in word finding, connected speech, reading and spelling, whereas auditory comprehension remained relatively stable. fMRI demonstrated progressively increasing activation within left-hemisphere language regions, reaching normative range by the final assessment, along with increasing activation in semantic/executive regions.

**Discussion:** Recovery of language production abilities can continue for years after stroke, supported by neuroplastic changes in the brain. Neurologists should inform stroke survivors that aphasia recovery can continue indefinitely after stroke.

## Introduction

Recovery from post-stroke aphasia has traditionally been thought to occur within the first 6-12 months after stroke, with improvement plateauing thereafter^1^. However, growing evidence suggests that later language improvement remains possible. Clinical trials demonstrate that speech-language therapy remains effective years after stroke^2^. A few longitudinal studies have reported continued group-level improvement over years^3,4^. However, averaging across participants may obscure individual changes. Moreover, these studies included only two timepoints and relied on only overall aphasia severity^3,4^, precluding insight into domain-specific language recovery.

Longitudinal neuroimaging studies within the first year after stroke ^5^, and cross-sectional studies in the chronic phase^6–8^, have revealed neuroplastic mechanisms supporting aphasia recovery. These studies implicate left-hemisphere language regions, right-hemisphere homologues, domain-general regions, and perilesional cortex as candidate substrates of recovery. However, to our knowledge, no study has longitudinally characterized changes in language activity over several years alongside detailed behavioral assessment.

We report a longitudinal case of chronic post-stroke aphasia in which language function improved across four timepoints spanning almost six years. This provides converging behavioral and neuroimaging evidence that aphasia recovery can continue for years after stroke.

## Materials and Methods

### Experimental Design

The case participant enrolled in two IRB-approved observational studies of chronic stroke outcomes at Georgetown University. Aphasia assessments and MRI sessions were performed at 9-, 22-, 33- and 71-months after stroke. Tests included the Western Aphasia Battery- Revised (WAB-R), and measures of auditory processing, executive functioning, phonology and semantics, naming, fluency, and motor speech (see Supplemental Table 1 for details).

MRI was acquired on a 3T Siemens Prisma scanner. Language activity was measured using an adaptive semantic matching fMRI paradigm, with a pseudofont matching control condition^9^. 116 healthy older controls completed the same fMRI paradigm (described in Anderson et al., 2026; Supplemental 2)^10^.

Mean contrast estimates (semantic>pseudofont blocks) were extracted from five regions of interest (ROIs): left language, left semantic/executive function, right language, right semantic/executive function, and perilesional^8^. Data from 20 controls were used to define ROIs. Mean ROI contrast estimates from the case participant were compared with the remaining 96 healthy controls^10^. To test for change across the four timepoints, a first-level omnibus F-contrast tested the Task×Session interaction. Significant interactions were interrogated using post-hoc t-contrasts testing for increases in activation as a function of log(months post-stroke) within each ROI, and Bonferroni corrected.

Longitudinal voxel-wise changes in activation were modeled as a function of log(months post-stroke), thresholded using voxel-level family-wise error correction^11^.

### Standard Protocol Approvals, Registrations, and Patient Consents

The participants provided written informed consent under Georgetown University IRB-approved protocols. The case participant reviewed the manuscript and signed a consent-to-disclose. Data not published within this article are not shared.

## Results

The case participant is a 46-year-old right-handed lawyer with no known past medical history who presented with acute right hemiparesis and aphasia, treated with intravenous alteplase (tPA/Activase) and successful mechanical thrombectomy. Examination revealed right-sided hemiparesis, neglect and absent withdrawal to pain. Initial National Institutes of Health Stroke Scale score was 21. CT angiography demonstrated occlusion of left internal carotid artery and distal left middle cerebral artery. MRI showed an ischemic stroke of superior temporal, posterior frontal, inferior parietal, and insular cortices, and the putamen. The WAB-R classification was Broca’s aphasia. Speech was nonfluent and agrammatic, characterized by short utterances and frequent paraphasia, impaired auditory comprehension, and reliance on orthographic cueing to support comprehension and word finding. On an in-house therapy engagement survey, he reported practicing language “nearly every day” since his stroke through structured home exercises, reading with audiobook support, language applications and frequent daily conversation (6+ hours/day).

WAB-R Aphasia Quotient (AQ) increased from 49.4 at 9 months to 65.2 at 22 months and 71.5 at 33 months, before declining slightly to 69.2 at 71 months (Figure 1A). Improvements were observed in spontaneous speech, word finding, picture description, naming, category fluency, oral reading, and spelling, whereas auditory verbal comprehension and executive function remained relatively stable. Real word reading improved progressively, whereas pseudoword reading remained impaired throughout follow-up (Figure 1; Supplemental Table 1). Motor speech showed early improvement followed by stabilization.

**Figure 1.**
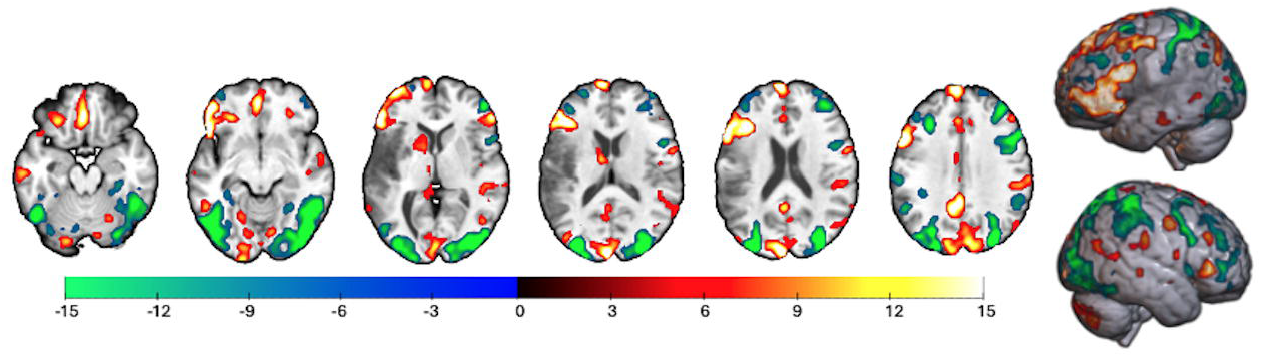
Behavioral performance improved across multiple language domains. Behavioral performance across selected assessments obtained 9, 22, 33, and 71 months after stroke. (A) Western Aphasia Battery-Revised (WAB-R) Aphasia Quotient (AQ) and subscale scores. WAB AQ increased over time, with improvements in word finding and spontaneous speech, whereas auditory verbal comprehension and repetition remained relatively stable. (B) WAB picture description measures, measured by content units per minute, mean length of utterance, words per minute, and total words produced. Picture description improved across all measures. (C) Picture and verb naming accuracy. Naming accuracy improved overall with larger gains in picture naming than verb naming. (D) Oral real word reading accuracy increases steadily throughout follow-up. (E) Spelling measured by a typing-to-dictation task and quantified as 1-Levenstein distance, where values closer to 1 indicate greater similarity between the typed response and target word. (F) Real-word and pseudoword repetition accuracy remain relatively stable, whereas pseudoword repetition showed modest improvement and remained impaired.

Longitudinal fMRI demonstrated significant changes in activation across the four timepoints in five ROIs (left language: 15.1 =, p = 5.01 × 10^−09^; left semantic: F = 16.5, p = 6.70 × 10^−10^; right language: F = 6.33, p = 1.4 × 10^−3^; right semantic: F = 6.75, p = 7.88 × 10^−4^; perilesional: F = 9.13, p = 2.65 × 10^−5^; Figure 2; Supplemental Table 2). Left language ROI activity followed an approximate logarithmic curve with activity increasing to near the control mean level by the final assessment. Activity in the semantic/executive ROI increased to above the control range by the third timepoint, with a slight decline at the final assessment. Right language and semantic/executive ROI activity increased linearly from the first to the third timepoint and then decreased to the control mean level at the fourth timepoint. Perilesional ROI activity increased to the control mean at the third timepoint and remained stable at the fourth. The voxelwise analysis demonstrated large clusters of increased activity in residual left frontal language regions adjacent to the lesion, with patchy areas of increased activation elsewhere, and decreased activation in bilateral frontoparietal and occipital regions (Figure 3).

**Figure 2.**
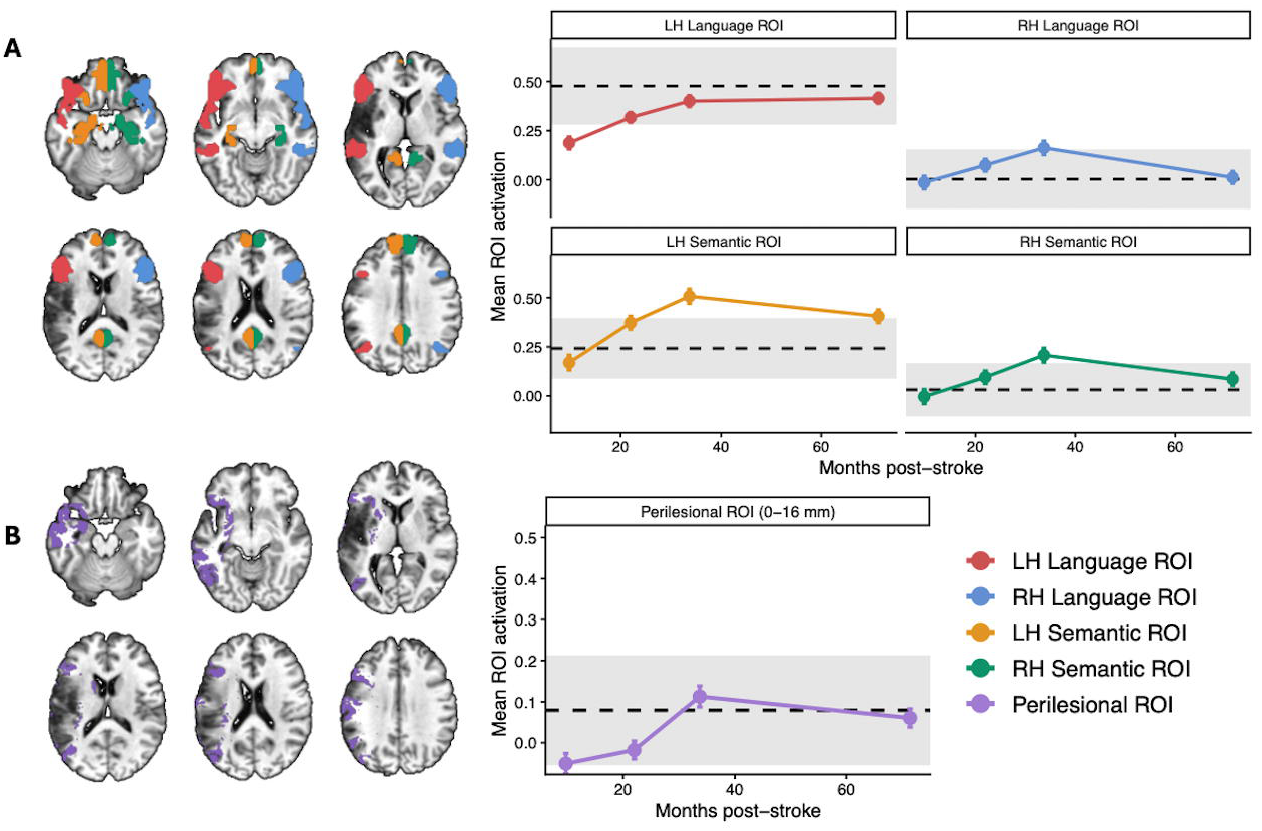
Longitudinal changes in language-related fMRI activation. Left panels show regions of interest (ROIs), and the right panels show BXC’s mean task-related activation (semantic > control contrast beta values) within each ROI across the four imaging sessions. Points indicate BXC’s mean ROI activation at each timepoint. Error bars represent the within-subject standard error of the contrast estimate, derived from the first-level GLM. The dashed line indicates the mean activation of the healthy control group (n = 96), and the grey shaded region represents +/−1 between-subject standard deviation of the control group mean ROI activation. (A) Functionally defined language (red, blue) and semantic/executive (orange, green) regions of interest. Language and Semantic ROIs were defined from an independent cohort of 20 healthy controls who completed the same task, and homologous right hemisphere ROIs were generated by mirroring the left hemisphere ROIs across the midline. The same ROIs were used and described in Turkeltaub et al (2025). (B) Perilesional ROI (purple) generated by dilating the lesion mask by 16 mm and intersecting the resulting mask with a gray matter mask.

**Figure 3.**
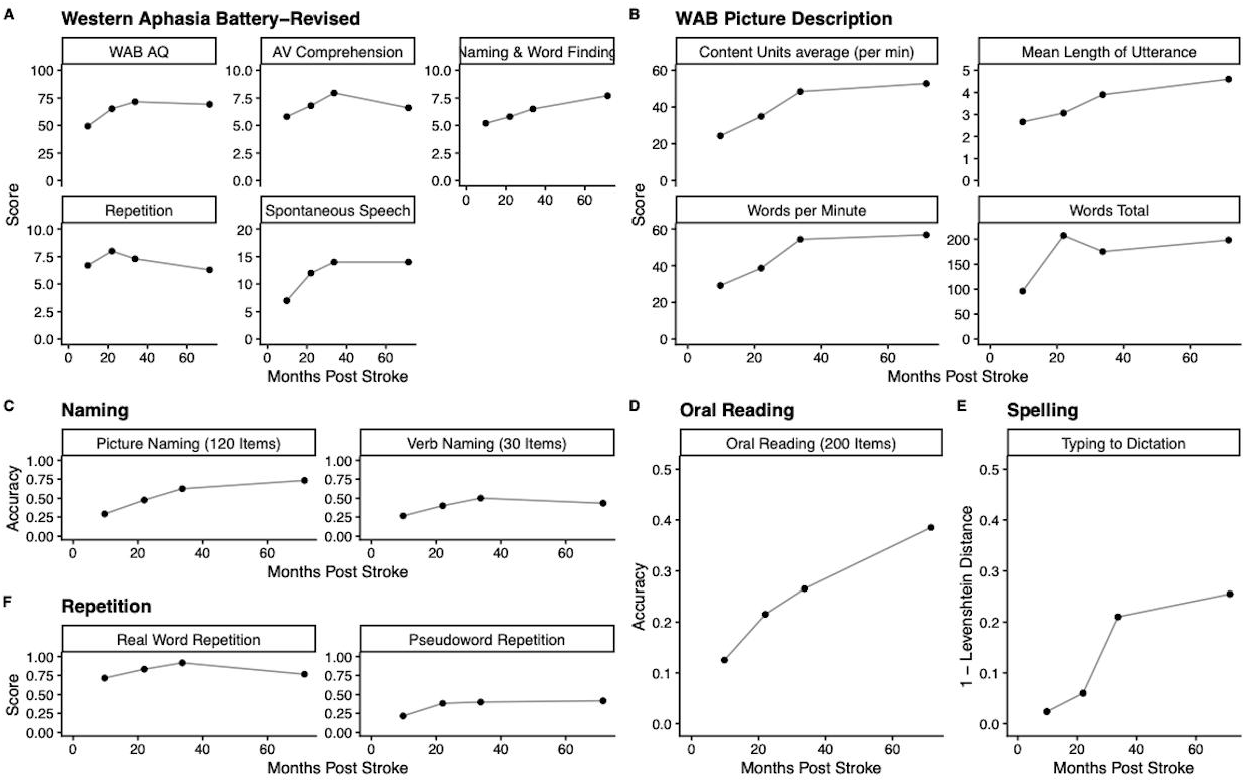
Voxelwise longitudinal changes in semantic decision fMRI activation. Whole-brain analysis identified regions in which activation changed as a function of log(months post-stroke). Warm colors indicate increasing activation over time and cool colors indicate decreasing activation. Statistical maps are thresholded at voxel-level FWE-corrected p < 0.05 and cluster size ≥30 voxels.

## Discussion

This case demonstrates that aphasia recovery can continue for years after stroke. Over six years, the case participant improved across multiple language domains, extending previous reports of continued recovery. The nearly 20-point improvement in WAB-R AQ substantially exceeded the approximately 5-point change commonly considered clinically meaningful in aphasia rehabilitation studies^12^. Importantly, recovery was not uniform across language functions. Lexical retrieval, reading, spelling, and connected speech improved substantially, with ongoing improvement even several years after stroke, whereas auditory comprehension and repetition improved over the first few years, but modestly worsened later. These findings suggest that chronic aphasia recovery is domain-specific rather than reflecting a generalized improvement. Behavioral trajectories paralleled changes in different functional ROIs, suggesting that activation patterns may relate to recovery of specific language functions, although this requires further investigation.

Recovery after left-hemisphere stroke has been hypothesized to be supported by several neuroplastic mechanisms, including recruitment of perilesional cortex^7,13^, normalization of residual left-hemisphere language networks^6^, compensatory right-hemisphere activation^8^, and engagement of domain-general cognitive control regions^14^. Evidence supporting these models come from studies of acute recovery or cross-sectional analyses of chronic aphasia, leaving their relative contributions during long-term recovery unclear. Neuroplastic mechanisms underlying aphasia recovery likely vary across stroke survivors, so examining individual cases is essential^15^. The largest longitudinal change occurred within the left-hemisphere language regions, where activation increased from below the normative range to values approaching those of healthy controls. Perilesional activation also increased but did not exceed the normative range, suggesting normalization of activity within residual language cortex rather than the emergence of novel compensatory activation^6^. This pattern is consistent with the case participant, relying primarily on restoration of function within the residual left-hemisphere language network, and possibly compensation in domain-general regions, whereas right-hemisphere compensation may have peaked a few years after stroke and then declined.

The case participant sustained engagement in speech-language therapy and independent language practice may have contributed to the prolonged improvements observed. These findings complement evidence from randomized clinical trials demonstrating that speech-language therapy remains effective during the chronic phase of aphasia. This case illustrates that chronic aphasia recovery is dynamic, domain-specific, and accompanied by evolving brain activation. Larger longitudinal studies integrating comprehensive behavioral assessment with functional neuroimaging will be needed to identify continued recovery and determine which patients benefit from different rehabilitation approaches long after stroke. In conclusion, neurologists should not tell stroke survivors that aphasia recovery ends after one year, but rather that, for some, recovery can continue for many years, supported by ongoing neuroplasticity.

