## Supplement 2 for "Dispelling the Plateau Assumption: Case-Based Evidence for Language Recovery in Chronic-Post Stroke Aphasia"

|  |  | LH Language | LH Semantic | RH Language | RH Semantic | Perilesional |
| --- | --- | --- | --- | --- | --- | --- |
| ANOVA | F | 15.12 | 16.52 | 6.33 | 6.75 | 9.13 |
|  | p | 1.00E-09* | 1.34E-10* | 0.00* | 0.00* | 5.31E-06* |
|  | df | 0.83, 1810 | 0.83, 1810 | 0.83, 1810 | 0.83, 1810 | 0.83, 1810 |
| T2 vs T1 | F | 12.65 | 18.56 | 4.51 | 4.79 | 1.01 |
|  | P | 0.00* | 1.73E-05* | 0.03* | 0.03* | 0.32 |
| T3 vs T2 | F | 5.80 | 8.60 | 4.19 | 6.32 | 15.04 |
|  | P | 0.02* | 0.00* | 0.04* | 0.01* | 0.00* |
| T4 vs T3 | F | 0.17 | 4.96 | 12.29 | 7.53 | 2.37 |
|  | P | 0.68 | 0.03* | 0.00* | 0.01* | 0.12 |
| T4 vs T1 | F | 37.97 | 25.40 | 0.39 | 3.92 | 11.08 |
|  | P | 8.71E-10* | 5.08E-07* | 0.53 | 0.05* | 0.00* |
| log(months) | t | 6.50 | 5.71 | 1.11 | 2.54 | 4.10 |
|  | p | 2.56E-10* | 3.35E-08* | 0.67 | 0.03* | 0.00* |

**Supplemental Table 2.** Omnibus F-contrasts tested whether task-related activation (words > control) differed across the four imaging sessions. Post-hoc two-sided t-contrasts tested pairwise differences between consecutive sessions and between the first and final sessions. The log(months) t-contrast tested for increasing activation as a function of log-transformed months post-stroke, with p-values Bonferroni-corrected across the five ROIs. Degrees of freedom are effective degrees of freedom estimated from the first-level fMRI model.

**Neuroimaging Data Acquisition and Pipeline**

Structural and functional MRI data were processed using the pipeline described by Anderson and colleagues (2026). Participants completed an adaptive semantic decision fMRI task to localize language-sensitive cortex. During the semantic condition, participants viewed pairs of words and indicated whether they were semantically related (e.g., calendar-date) using a button press. During the control condition, participants viewed pairs of pseudofonts and indicated whether they were visually identical. The task consisted of ten alternating blocks of semantic and control conditions. An adaptive staircase procedure adjusted task difficulty independently for the semantic and control conditions to maintain comparable performance across participants. Following two consecutive correct responses, task difficulty increased by one level; following an incorrect response, difficulty decreased by two levels. Difficulty was manipulated by varying lexical frequency, age of acquisition, word length, semantic relatedness, concreteness, and presentation rate for the semantic condition, and visual similarity of mismatching pseudofont pairs for the control condition. Presentation rate was adjusted at the beginning of each semantic block and held constant for the subsequent control block to equate task demands.

MRI data were acquired on a 3T Siemens MAGNETOM Prisma Fit scanner using a 20-channel head coil. Structural imaging included a high-resolution T1-weighted magnetization prepared rapid gradient echo (MPRAGE) and fluid-attenuated inversion recovery (FLAIR) sequence. First, a T1-weighted MPRAGE sequence was acquired: 176 sagittal slices; slice thickness = 1 mm; 1 mm^3^ voxels; field of view (FOV) = 256 mm; matrix = 256 x 256 mm; flip angle = 9º; generalized autocalibrating partial parallel acquisition = 2; repetition time (TR) = 1900 ms; echo time (TE) = 2.98 ms; scan time: ~ 5 mins. Second, a FLAIR sequence was acquired: 192 sagittal slices; slice thickness = 1 mm; 1 mm^3^ voxels; FOV = 256 mm; matrix = 256 x 256 mm; flip angle = 120º; generalized autocalibrating partial parallel acquisition = 2; TR = 5000 ms; TE = 386 ms; scan time: ~ 6 mins. To assist with registering the BOLD data to the anatomical, a coplanar T2-weighted image was also acquired: 48 slices; TR = 4010 ms; TE = 76 ms; slice thickness = 2.6mm with a 10% gap; 2.9 mm voxels; FA 150°.

To assess language activation, 504 volumes of a BOLD T2*-weighted scan were acquired: 48 slices; TR = 794 ms; TE = 30 ms; slice thickness = 2.6 mm with a 10% gap; 2.9 mm voxels; FOV= 211 mm; matrix = 74 x 74; FA 50°; SMS = 4; scan time ~ 7 mins. Spin echo field maps were acquired to correct magnetic field distortions: 48 slices; TR = 790 ms; TE = 30 ms; slice thickness = 2.6 mm with 10% gap; FOV = 211; matrix = 74x74; FA = 90°; scan time: ~7 mins.

*Pipeline*

Functional MRI preprocessing was performed using SPM12 (https://www.fil.ion.ucl.ac.uk/spm/software/spm12/). Preprocessing included motion correction, coregistration to the participant's anatomical image using the coplanar T2-weighted scan, spatial normalization to MNI152 space (using Advanced Normalization Tools, <https://picsl.upenn.edu/software/ants/>), and spatial smoothing with a 6-mm FWHM Gaussian kernel. The location of the stroke lesion was manually segmented and applied as a cost-function mask during spatial normalization.

A first-level fixed-effects general linear model was estimated separately for each imaging session. The model included regressors for the semantic decision task and the perceptual control task, along with 12 nuisance regressors consisting of the six head-motion parameters and their first temporal derivatives. Task effects were modeled using the canonical hemodynamic response function, and the contrast of semantic decision greater than perceptual control (words > control) was computed for each session, producing voxelwise contrast images for subsequent whole-brain and region-of-interest analyses.

Longitudinal changes in activation were assessed using SPM. An omnibus F-contrast tested for a Task × Session interaction across the four imaging sessions. Significant omnibus effects were interrogated using post hoc t-contrasts testing for increases in activation as a function of log-transformed months post-stroke. Whole-brain analyses were thresholded using voxel-level family-wise error correction (p < 0.05).

*Regions of Interest*

Mean activation was extracted using the MarsBar Toolbox (<http://marsbar.sourceforge.net/>) from five predefined regions of interest: left language, right language, left semantic/executive, right semantic/executive, and a perilesional gray matter ROI extending 0–16 mm from the lesion boundary. Language and semantic/executive ROIs were defined independently from task activation in healthy controls, with homologous right hemisphere ROIs generated by mirroring across the midline. The perilesional ROI was generated by dilating the manually segmented lesion and intersecting the resulting shell with gray matter as determined by values > 0.5 in the grey matter SPM T1 segmentation image. Mean contrast estimates and voxelwise standard deviation estimates were extracted from each ROI for each imaging session. Statistical significance for ROI analyses was Bonferroni corrected across the five ROIs.
