## Supplement 1 for "Dispelling the Plateau Assumption: Case-Based Evidence for Language Recovery in Chronic-Post Stroke Aphasia"

| **Measure Name** | **First** | **Second** | **Third** | **Fourth** | **Task Description** |
| --- | --- | --- | --- | --- | --- |
| **Auditory Perception and Comprehension** | | | | |  |
| Audiovisual word-to-text matching accuracy | 0.66 | 0.66 | 0.71 | 0.77 | 40 item test. The participant sees a video of a person producing the target word. On the next screen, the participant is prompted to select the written target from a field of four. |
| Auditory word-to-text matching accuracy | 0.54 | 0.61 | 0.59 | 0.69 | 40 item test. The participant is presented the target word via headphones. On the next screen, the participant is prompted to select the written target from a field of 4. |
| Environmental sound-to-text matching accuracy | 0.9 | 0.92 | 0.98 | 0.95 | 40 item test. The participant is presented the environmental sound target via headphones. On the next screen, the participant is prompted to select the written target from a field of 4. |
| Auditory word-to-picture matching accuracy | 0.55 | 0.65 | 0.9 | 0.79 | See (Lacey et al., 2017) |
| NAVS sentence comprehension accuracy (Canonical) | 0.67 | 0.57 | 0.47 | 0.67 | See (Cho-Reyes & Thompson, 2012) |
| NAVS sentence comprehension accuracy (Non-Canonical) | 0.67 | 0.67 | 0.67 | 0.47 | See (Cho-Reyes & Thompson, 2012) |
| **Executive function/Working Memory** | | | | |  |
| Antelopes and cantaloupes standard (Pattern Length 1) | 20 | 21 | 23 | 23.7 | See (McCall et al., 2022) |
| Antelopes and cantaloupes standard (Pattern Length 2) | 18 | 18.5 | 21.5 | 18.7 | See (McCall et al., 2022) |
| Digit span backward total span score | 0 | 3 | 2 | 1 | Sequences of 2 up to 9 |
| Digit span forward total span score | 4 | 4 | 4 | 4 | Sequences of 2 up to 9 |
| **Fluency** | | | | |  |
| Category Fluency (Animals, Fruits and Vegetables Average) | 4 | 5.5 | 6.5 | 10.5 |  |
| Letter Fluency (F,A,S Average) | 1 | 3.7 | 5.7 | 3.7 |  |
| **General aphasia severity** | | | | |  |
| WAB-R Aphasia Quotient | 49.4 | 65.2 | 71.5 | 69.2 | See (Kertesz, 2007) |
| WAB-R AV Comp | 5.8 | 6.8 | 7.95 | 6.6 | See (Kertesz, 2007) |
| WAB-R Word Finding and Naming | 5.2 | 5.8 | 6.5 | 7.7 | See (Kertesz, 2007) |
| WAB-R Repetition | 6.7 | 8 | 7.3 | 6.3 | See (Kertesz, 2007) |
| WAB-R Spontaneous Speech | 7 | 12 | 14 | 14 | See (Kertesz, 2007) |
| **Motor speech** | | | | |  |
| Apraxia of Speech Rating Scale Total Rating | 18 | 11 | 11 | 11 | See (Duffy et al., 2023) |
| Motor speech exam alternating motion rate (per second) | 0.80 | 3.47 | 4.07 | 4.27 | See (Haley et al., 2012) |
| Motor speech exam sequential motion rates (per second) | 0.2 | 0.6 | 0 | 0 | See (Haley et al., 2012) |
| **Naming** | | | | |  |
| Picture Naming Accuracy | 0.29 | 0.48 | 0.62 | 0.73 | See (Fama et al., 2019) |
| Verb Naming Accuracy | 0.27 | 0.4 | 0.5 | 0.43 | 30 item action picture naming test using line drawings. |
| **Phonology** | | | | |  |
| Pseudoword repetition accuracy | 0.22 | 0.38 | 0.4 | 0.42 | See (Fama et al., 2019) |
| Real word repetition accuracy | 0.72 | 0.83 | 0.92 | 0.77 | See (Fama et al., 2019) |
| Rhyme judgement auditory accuracy | 0.55 | 0.68 | 0.85 | 0.8 | See (Fama et al., 2019) |
| Rhyme production accuracy | 0.04 | 0 | 0.08 | 0.24 | 25 item test with words presented via headphones. The participant is prompted to produce a word that rhymes with the target. |
| TALSA phoneme discriminations on pseudowords accuracy | 0.95 | 0.8 | 0.89 | 0.77 | See (Martin et al., 2018) |
| **Reading** | | | | |  |
| Oral reading accuracy | 0.12 | 0.22 | 0.26 | 0.38 | See (Anderson et al., 2026; Dickens et al., 2021) |
| Pseudoword reading accuracy | 0.02 | 0.02 | 0.02 | 0 | See (Anderson et al., 2026; Dyslin et al., 2025) |
| Written lexical decision accuracy | 0.78 | 0.76 | 0.77 | 0.73 | See (Anderson et al., 2026; Chang et al., 2026) |
| **Semantic** | | | | |  |
| Pyramids and Palm Trees accuracy | 0.88 | 0.92 | 0.9 | 0.93 | See (Howard & Patterson, 1992) |
| TALSA category judgement picture accuracy | 0.93 | 0.92 | 0.98 | 0.98 | See (Martin et al., 2018) |
| **Spelling / typing** | | | | |  |
| Typing-to-dictation accuracy | 0 | 0.09 | 0.09 | 0.19 | 32 item test with words presented via headphones. The participant is prompted to spell the target. |
| Typing-to-dictation Levenstein distance | 0.98 | 0.94 | 0.79 | 0.75 | 32 item test with words presented via headphones. The participant is prompted to spell the target. |
